# Single-cell hit calling in high-content imaging screens with Buscar

**DOI:** 10.64898/2026.04.15.718737

**Authors:** Erik Serrano, Wei-shan Li, Gregory Way

## Abstract

High-content screening (HCS) enables the systematic quantification of single-cell morphology features across thousands of perturbations, capturing rich phenotypic heterogeneity. Image-based profiling is a critical bioinformatics processing step in this pipeline, as researchers use it to predict mechanisms of action, assess toxicity, perform hit calling, and more. However, current image-based profiling workflows rely on aggregate statistics, such as calculating mean or median feature values per well, implicitly assuming cell homogeneity. This limitation obscures subpopulation effects, reducing sensitivity to subtle or heterogeneous effects of perturbations. Here we present Buscar, a method that leverages the full heterogeneity of single-cell image-based profiles to call hits. Buscar requires two reference, single-cell populations that define distinct morphology states: a reference state (e.g., disease cells) and a target state (e.g., healthy cells). Buscar then compares these two groups to define on- and off-morphology signatures, which it then uses to score every perturbation in a given screen. The scores quantify perturbation efficacy and off-target effects, or specificity, in an interpretable manner, clarifying which morphologies are appropriately altered and which may arise from off-target activity. We apply Buscar to three datasets. First, as a proof of concept, we applied Buscar to a Cell Painting dataset of cardiac fibroblasts from patients with heart failure. Buscar quantifies both morphology rescue and off-target morphology activity in these cells treated with a TGFβ receptor inhibitor. Second, we show that Buscar recovers biologically coherent gene-phenotype associations across 16 manually-labeled phenotypes in the MitoCheck dataset. Lastly, applied to CPJUMP1, we show that Buscar is robust to technical replicates collected across plates in both small-molecule and CRISPR-Cas9 perturbations. Together, these results establish Buscar as a reproducible and interpretable hit calling method that overcomes aggregation bias, enabling the simultaneous quantification of compound efficacy and specificity to enhance hit calling in HCS. We release Buscar as an open-source python package.

## Introduction

High-content screening (HCS) integrates automated microscopy with sophisticated image analysis to systematically measure the effects of perturbations on cells in a high-throughput manner (Caicedo *et al*., 2017; Way *et al*., 2023; Lin *et al*., 2020; Seal *et al*., 2025; Krentzel *et al*., 2023; Serrano *et al*., 2026). This approach has revolutionized phenotypic drug discovery by providing insights into disease mechanisms and identifying new therapeutic agents (Way *et al*., 2023; Vincent *et al*., 2022). HCS begins with cells cultured in multi-well plates and exposed to chemical compounds or genetic perturbations, followed by multiplex fluorescence staining. This is most commonly done using the Cell Painting assay, which employs six dyes imaged across five channels to visualize eight cell components and organelles: DNA, cytoplasmic RNA, nucleoli, actin, Golgi apparatus, plasma membrane, endoplasmic reticulum, and mitochondria (Bray *et al*., 2016). An automated microscope then captures images of the marked cells across all experimental conditions. These images undergo a computational processing pipeline that includes illumination correction and quality control to reduce technical artifacts, followed by segmentation to identify individual cell objects within the images (Caicedo *et al*., 2017). Feature extraction then quantifies hundreds to thousands of morphological descriptors per cell, spanning shape, size, intensity, texture, and granularity across all fluorescent channels, using open-source tools such as CellProfiler (Stirling *et al*., 2021) and CHAMMI (Agrawal *et al*., 2026), or commercially available platforms like those from Revvity and IN Carta by Molecular Devices. The resulting image-based profiles then pass through a standardized computational workflow to ensure data quality and analytical readiness (Serrano *et al*., 2026; Driscoll and Zaritsky, 2021; Caicedo *et al*., 2017). First, researchers apply data harmonization to unify outputs into a schema-consistent dataset by resolving incompatible file formats, inconsistent column naming, and ambiguous data types, thus eliminating the need for custom integration code (Bunten *et al*., 2025). Then, researchers apply single-cell quality control (QC) to flag profiles that contain missegmented cells (whether over- or under-segmented), as well as debris and other contaminants (Tomkinson *et al*., 2025). Finally, researchers process image-based profiles (by applying annotation, normalization, feature selection, and batch effect correction) to prepare them for downstream analysis (Serrano *et al*., 2025). Together, these bioinformatics steps yield a rich, analysis-ready dataset in which each cell is represented as a high-dimensional feature vector encoding its morphology state.

Single-cell image-based profiles provide the foundation for downstream analyses aimed at characterizing phenotypic signatures associated with specific perturbations. The current standard practice, however, involves aggregating all features across single cells using summary statistics such as mean or median values, or alternatively using deep learning approaches to derive latent variable representations to characterize profiles at specific levels such as treatment or well (Caicedo *et al*., 2017; Tang *et al*., 2024). This is a critical limitation. Aggregation implicitly assumes that a given perturbation elicits a uniform phenotypic response across all cells, which may not reflect reality (van Dijk *et al*., 2024; Stossi *et al*., 2023; Wagner *et al*., 2025; Pearson *et al*., 2022). At minimum, collapsing single-cell measurements into summary statistics results in substantial loss of information, including cell-to-cell heterogeneity, the presence of distinct subpopulations (e.g. resistant or non-resistant cells), differential toxicity responses, and other phenotypic variations that are crucial for understanding perturbation mechanisms (Stossi *et al*., 2023; Gough *et al*., 2017; Maleki *et al*., 2024). Consequently, the failure to capture this heterogeneity may contribute to the high attrition rates observed in clinical development, where compounds that appeared promising in initial screens fail to demonstrate efficacy or safety in more complex biological contexts (Sun *et al*., 2022). It is therefore critical to develop analytical methods that accurately account for the biological nuances present in single-cell image-based profiles, thereby improving hit identification and reducing false positive rates in drug discovery pipelines.

Despite this limitation, researchers have made important discoveries and substantial progress in analyzing aggregated image-based profiles. The standard analysis calculates high-dimensional distance metrics such as cosine similarity, Mahalanobis distance (Vulliard *et al*., 2022; Hutz *et al*., 2013; Nyffeler *et al*., 2021), Pearson correlation, or Euclidean distance (Kalinin *et al*., 2025). Using this approach, researchers have successfully grouped compounds based on phenotypic similarities, a computational strategy commonly known as “guilt by association.” For example, multiple studies have demonstrated that compounds sharing common molecular targets display similar profiles irrespective of their chemical structures (Ziegler *et al*., 2021; Simm *et al*., 2018; Odje *et al*., 2024; Ringers *et al*., 2025). In addition, using supervised machine learning aggregated profiles have been used to predict the mechanism of action (MoA) of compounds (Pahl *et al*., 2023; Papanastasiou *et al*., 2025; Cox *et al*., 2020) and to identify on-target polypharmacology, revealing that effects often depend on engaging multiple molecular targets (Hughes *et al*., 2021; Hafner *et al*., 2019; Chow *et al*., 2022). These advances have collectively accelerated the identification of novel compounds for drug discovery (Gibson *et al*., 2015; Lo Cicero *et al*., 2016; Boyd *et al*., 2020). Beyond profile similarity, hit calling has its own dedicated methods using aggregated profiles. The Z’-factor (Zhang *et al*., 1999) provides a widely used univariate metric for assay quality and hit identification, quantifying the separation between positive and negative control distributions in a single readout. Another approach utilized deep learning embeddings from bulk fluorescence miscopy images, projecting compound responses onto a perturbation vector between unperturbed and perturbed cell states to track rescue efficacy and morphological specificity (Cuccarese *et al*., 2020; Heiser *et al*., 2020). More recently, the mean average precision (mAP) framework (Kalinin *et al*., 2025) extended hit calling to high-dimensional aggregated profiles by assessing whether perturbation replicates are more similar to each other than to controls using ranked distance metrics. However, all three approaches operate on aggregated profiles, failing to account for the cellular heterogeneity present within treatment groups and neither quantifying perturbation efficacy nor morphological specificity at the single-cell level.

Here we introduce Buscar (Bioactive Unbiased Single-cell Compound Assessment and Ranking; Spanish for “*to look for*”), a method for reproducible hit-calling that operates directly on distributions of single-cell profiles. Buscar requires two reference populations defining distinct morphology states (e.g., diseased and healthy cells). This is essential because it enables Buscar to separate the high-dimensional feature space into two complementary, mutually exclusive signatures: an on-morphology signature (features that differ significantly between the reference and target states) and an off-morphology signature (features that remain unchanged).

This separation allows the independent tracking of efficacy and specificity per perturbation in the screen. In contrast, conventional methods relying on aggregated profiles and only one reference population lose single-cell heterogeneity and cannot distinguish efficacy from specificity. Next, Buscar uses these two signatures to compute two complementary metrics for each perturbation: an on-Buscar score, which quantifies efficacy, and an off-Buscar score, which quantifies specificity. The on-Buscar score measures distributional similarity by computing Earth Mover’s Distance (EMD) (Orlova *et al*., 2016) using the on-morphology signature features, while the off-Buscar score captures off-target effects by quantifying the extent to which off-morphology signature features are significantly altered under perturbation. By retaining cell-to-cell variability, Buscar captures phenotypic heterogeneity, including subpopulation structure and differential responses, that are typically obscured by aggregation. As a proof of concept, we apply Buscar to a dataset of failing and healthy cardiac fibroblasts. We demonstrate that it detects phenotypic rescue (high efficacy/low on-Buscar scores) while simultaneously identifying subtle off-target effects (captured by the off-Buscar score) associated with TGFβ receptor inhibition (Travers *et al*., 2025). We further demonstrate that Buscar recovers phenotype-specific gene rankings consistent with known mitotic and nuclear morphology biology in the MitoCheck dataset (Neumann *et al*., 2010) and achieves robust cross-plate reproducibility across both small-molecule and CRISPR–Cas9 perturbations in the CPJUMP1 dataset (Chandrasekaran *et al*., 2024). Together, these results establish Buscar as an interpretable and reproducible method for single-cell hit calling.

## Results

### Introducing the Buscar method

Buscar is a reproducible computational method and Python package that scores perturbations in high-content screens based on their efficacy and specificity using single-cell, image-based profiles (**Figure 1**). Buscar operates through two sequential modules: (1) Defining morphology signatures, and (2) Perturbation efficacy and specificity scoring.

**Figure 1.**
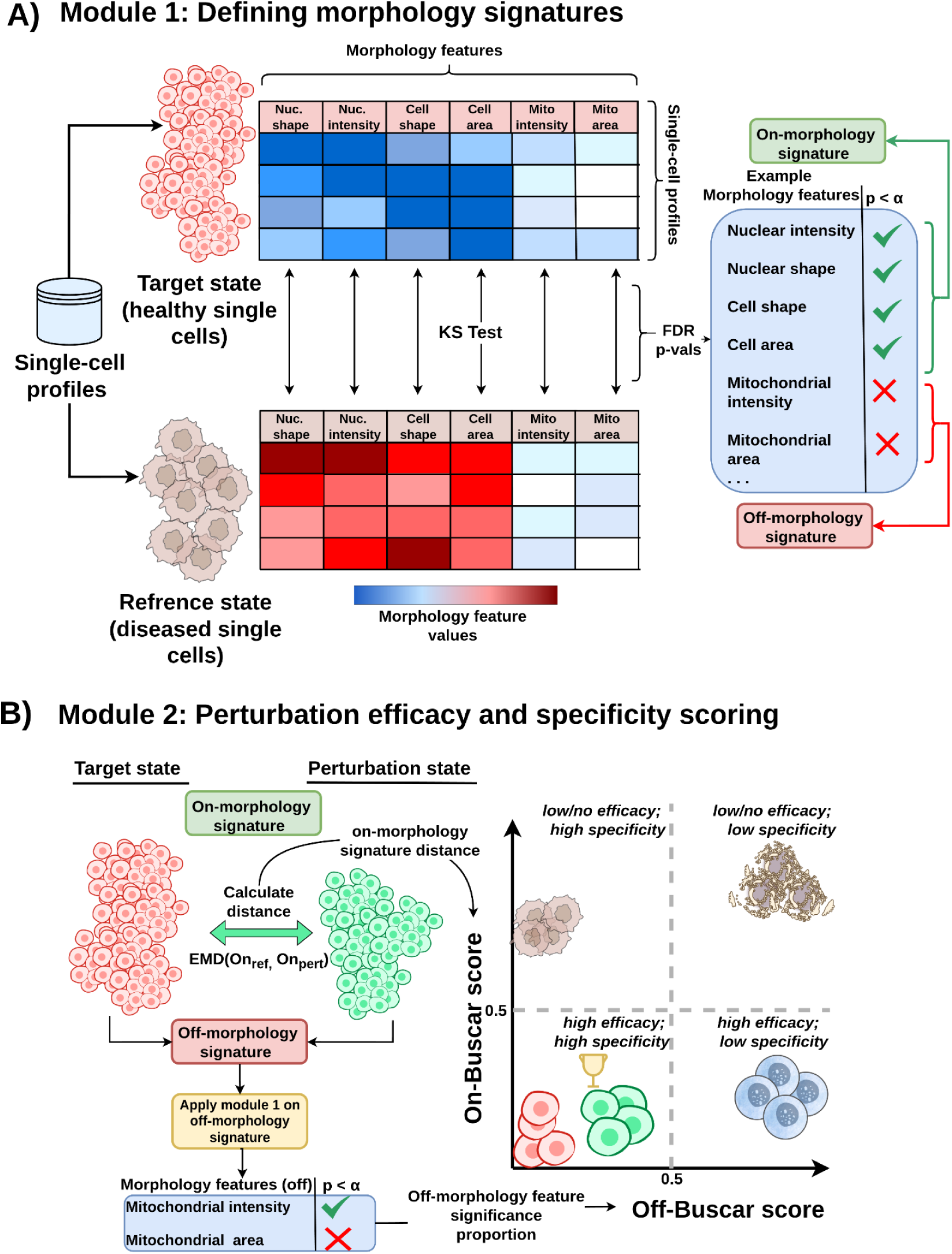
The Buscar method for single-cell hit calling. **(A)** Buscar defines on- and off-morphology signatures by identifying morphological features that distinguish, or do not distinguish, a negative control reference (e.g., disease cells) from a target state (e.g., healthy cells). **(B)** Buscar then uses these signatures to generate an on-Buscar score quantifying perturbation efficacy and an off-Buscar score quantifying specificity (i.e., off-target effects). Dashed lines indicate the 0.5 threshold on both axes and are shown for visual reference only.

#### Module 1: Defining morphology signatures (***Figure 1A***)

The first module establishes the morphology reference for evaluating perturbations, requiring two control populations that define distinct morphology states: a reference state and a target state. In practice, the reference state is a negative control (e.g., DMSO-treated diseased cells), and the target state represents cells with a healthy phenotype. Buscar then applies a series of non-parametric statistical tests (e.g., Kolmogorov-Smirnov [KS] test) to each morphology feature comparing the two states. After correcting resulting p-values for multiple comparisons using False Discovery Rate (FDR), Buscar determines if the feature distributions differ significantly between the two states. Buscar groups features showing significant differences into an on-morphology signature, representing the set of morphology changes distinguishing the reference and target states. Additionally, Buscar groups features that do not differ significantly into an off-morphology signature, representing features expected to remain unchanged. Together, these signatures define the morphology basis for evaluating perturbations. Specifically, a desired perturbation will shift the distribution of cells from the reference state toward the target state along the on-morphology feature axis, while minimizing changes in the off-morphology feature axis. This enables Buscar to track efficacy and specificity for every perturbation.

#### Module 2: Perturbation efficacy and specificity scoring (***Figure 1B***)

The second module scores each perturbation in terms of efficacy and specificity by computing two complementary metrics: an on-Buscar score and an off-Buscar score. The on-Buscar score measures how closely the distribution of perturbed cells aligns with the target state along the on-morphology signature axis. Buscar computes this score by calculating EMD (Orlova *et al*., 2016; Rubner, 2000) between the perturbed and target single-cell populations using the on-morphology signature features, where a lower score indicates a close proximity of the perturbed single-cell population to the target population, reflecting higher efficacy. Buscar normalizes the on-Buscar score by dividing the perturbation EMD by the EMD calculated between target and reference. Furthermore, the off-Buscar score quantifies specificity of the perturbation, or off-target effects. It measures the proportion of off-morphology signature features that differ significantly between the perturbed and target single-cell populations. Since Buscar initially categorized these features as unaffected between the reference and target states, any change induced by the perturbation suggests an unintended, off-target or adverse effect. A lower off-Buscar score indicates fewer off-target effects and thus higher specificity. Together, these complementary scores provide a joint, informative method for compound prioritization, allowing users to visually assess and rank the balance of efficacy and specificity and select the best-performing perturbations for further study.

We release Buscar as an open-source Python package. We specifically designed Buscar to be compatible with the Cytomining Ecosystem, ensuring seamless interoperability with tools like Pycytominer, coSMicQC, and CytoTable. It leverages Polars for fast and scalable single-cell data processing and ensures reproducibility across computing environments via comprehensive unit testing and automated dependency checks. The package supports diverse perturbation types, including small molecules, genetic perturbations (e.g., CRISPR and RNAi), and other emerging modalities.

### Buscar reveals phenotypic signatures and rescue dynamics in cardiac fibroblasts

As a proof-of-concept evaluating Buscar, we applied it to a Cell Painting dataset of both healthy and failing cardiac fibroblasts (CFs) (Travers *et al*., 2025). The failing CFs were derived from patients with idiopathic dilated cardiomyopathy, a condition of heart failure characterized by extensive collagen accumulation in the myocardium as a consequence of chronic fibrosis (Travers *et al*., 2025; Schultheiss *et al*., 2019). Cardiac fibrosis is driven in part by aberrant TGFβ signaling, which promotes the activation of CFs into myofibroblasts, leading to excessive extracellular matrix deposition and progressive loss of cardiac function (Maruyama and Imanaka-Yoshida, 2022). In this dataset, failing CF cells were treated with a TGFβ receptor inhibitor (TGFβRi) tool compound, which is known to suppress fibrotic signaling (Park and Yoo, 2022; Shi *et al*., 2022; Parichatikanond *et al*., 2020). Given that TGFβ activity is elevated in fibrotic CFs, TGFβRi provides a biologically-relevant perturbation aimed at attenuating the fibrotic process and promoting a shift toward a healthier CF phenotype. Walking through each Buscar module to investigate how each behaves in a controlled use-case, we apply Buscar to single-cell profiles in this dataset **(Figure 2A)**.

**Figure 2:**
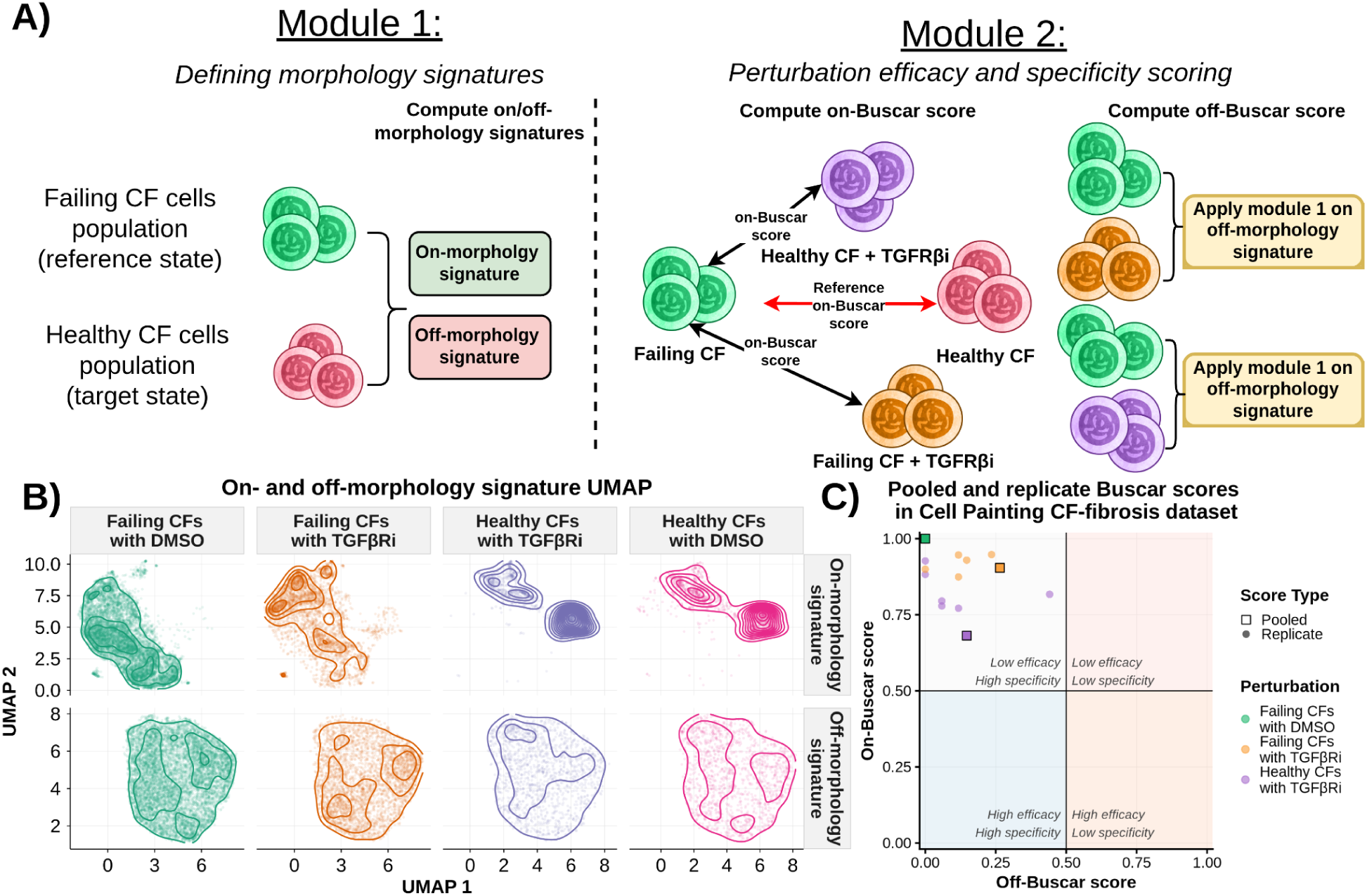
Applying Buscar to a Cell Painting dataset of healthy and failing cardiac fibroblasts (CFs). (**A)** Module 1 illustrates the generation of on- and off-morphology signatures from reference (failing) and target (healthy) CF states, while Module 2 leverages these signatures to calculate on- and off-Buscar scores to quantify perturbation efficacy and specificity. (**B)** UMAP projections of single-cell profiles computed independently on on-morphology (top row) and off-morphology (bottom row) signature feature, with kernel density estimate contours overlaid for each treatment condition: failing CFs with DMSO, failing CFs with TGFβRi, healthy CFs with TGFβRi, and healthy CFs with DMSO. (**C)** A scatter plot of on-Buscar scores (y-axis) versus off-Buscar scores (x-axis) for each treatment. Lines indicate the 0.5 threshold on both axes, dividing the plot into arbitrary quadrants for simplicity, each annotated to describe its corresponding region. Circles represent replicate treatment wells, whereas squares denote pooled populations (all single cells across wells that have the same treatment combined).

Using DMSO-treated failing and healthy CFs as the reference and target states, respectively, we first applied Buscar’s signature module (Module 1) to characterize the on- and off-morphology signatures (based on CellProfiler features). Examining the resulting signatures revealed that morphology changes between failing and healthy CFs were broadly distributed across all compartments, and channels indicating a global remodeling of cell morphology in the failing state **(Supplementary Figure 1**). When randomly shuffling failing and healthy labels, the Buscar signature module correctly allocated no feature to the on-morphology signature.

To visualize how these signatures structure the morphology landscape of each treatment, we projected all single-cell profiles into UMAP space using on- and off-morphology signatures separately (**Figure 2B**). In the on-morphology signature UMAP, failing DMSO and healthy DMSO cells occupied clearly distinct regions, confirming that the on-morphology signature captures relevant morphology differences between the healthy and failing states. In addition, the healthy DMSO population split into two distinct islands, revealing the presence of subpopulations with different phenotypes within the healthy CF population, a level of cell heterogeneity that would be entirely obscured by conventional aggregation approaches.

Because Buscar operates at single-cell resolution, both subpopulations are preserved and contribute to the scoring. Notably, the failing TGFβRi-treated cells shifted toward the healthy DMSO distribution, while healthy TGFβRi-treated cells remained clustered near the healthy DMSO population. In contrast, the off-morphology signature UMAP showed broadly overlapping distributions across all four conditions, demonstrating that off-features remain largely unchanged regardless of treatment, as expected.

Finally, we apply Buscar’s second module and calculate the on- and off-Buscar scores **(Figure 2C)**. Internally, Buscar normalizes the on-Buscar scores by the distance between the reference and target populations (i.e., healthy CFs treated with DMSO and failing CFs treated with DMSO). An on-Buscar score of 1.0 indicates a perturbation that is equivalent to the failing DMSO reference state, while an on-Buscar score of 0.0 would indicate a perturbation that induces a complete return to the healthy DMSO target state (**Supplementary Table 1**). When applied to healthy CFs treated with the TGFβRi tool compound, Buscar yielded an on-Buscar score of 0.68 and an off-Buscar score of 0.15. In contrast, applied to failing CFs treated with the same TGFβRi, the scores were higher, with an on-Buscar score of 0.90 and an off-Buscar score of 0.26. We also applied Buscar to individual replicate wells to evaluate score stability (**Supplementary Table 2**). We observed variation in replicate wells of healthy CFs treated with TGFβRi primarily along the on-Buscar score axis, which we observed to be negatively correlated with well-level cell counts (Spearman R = −0.86, p = 0.029; **Supplementary Figure 2A**), suggesting that EMD has sensitivity to distribution density in sparsely populated wells, which contributes to replicate variability in efficacy estimates. In contrast, replicate-well-level off-Buscar scores in healthy CFs treated with TGFβRi showed no significant correlation with cell count (Spearman R = 0.14, p = 0.785; **Supplementary Figure 2B**), suggesting that the off-Buscar scores are more robust to cell count variability. These results suggest that pooling cells across replicate wells, rather than scoring each well independently, may provide more stable on-Buscar score estimates. Looking at the pooled scores, the elevated off-Buscar score of failing CFs treated with TGFβRi compared to healthy CFs treated with TGFβRi suggests that TGFβRi perturbs additional morphology features specifically in the failing CF context, potentially reflecting secondary or compensatory responses to TGFβ pathway inhibition in the failing cell state. As well, TGFβRi does not fully reverse the failing CF phenotype indicating that a compound with a lower on-Buscar score would be a more promising candidate for follow-up testing for this disease with unmet clinical needs. Overall, this result demonstrates Buscar’s ability to simultaneously quantify compound efficacy and flag potential off-target effects, allowing users to rank perturbations in their ability to reverse disease phenotypes in much larger screens.

### Buscar scores are biologically-relevant across genes and phenotypes

To assess whether Buscar can prioritize biologically-relevant perturbations (which is a key requirement for ranking compounds in high-content screens), we analyzed gene-phenotype relationships in the MitoCheck dataset (Neumann *et al*., 2010). MitoCheck is a genome-wide RNAi screen targeting ∼21,000 human genes in HeLa cells expressing the chromatin marker H2B-GFP. Cells were imaged by time-lapse fluorescence microscopy and manually annotated into 16 phenotypes based on nuclear morphology. This process yielded a heterogeneous, single-cell mapping of individual phenotypes containing multiple gene perturbations and individual gene knockdowns spanning multiple phenotype classes. We can therefore test whether morphology profiles derived from genetic perturbations, regardless of phenotype label, retain gene-specific, biologically-relevant structure. Using feature-selected single-nucleus morphology features (Tomkinson *et al*., 2024), we performed a leave-one-gene-out (LOGO) Buscar analysis. Briefly, for each phenotype independently, we iteratively held out cells from each gene knockdown and used all non-held out cells of the remaining genes as the target population. In every iteration, we used interphase cells as the reference state, and applied Buscar. To assess significance, we used two permutation controls: (1) label permutation, in which gene labels were randomly shuffled 5,000 times to generate a null distribution, and (2) feature shuffling, in which morphological feature values were permuted column-wise to destroy the covariance structure while preserving label proportions. See methods for more details.

We observed that Buscar recovers gene-phenotype associations. For example, for phenotypes characterized by nuclear condensation including mitotic states (Neumann *et al*., 2010; Pau *et al*., 2013) and apoptosis (Frolova *et al*., 2023), gene perturbations with higher fractions of phenotype-labeled cells consistently received lower on- and off-Buscar scores (**Figure 3A**), indicating that genes whose knockdown yields a greater proportion of cells displaying the given nuclear condensation phenotype produce single-cell distributions that more closely resemble the phenotype-specific target population along the on-morphology signature axis, while inducing fewer changes along the off-morphology signature axis. We observed similar relationships with interphase-associated and multinucleated states, such as binuclear and polylobed phenotypes (Pau *et al*., 2013) (**Supplementary Figure 3**). In contrast, the feature shuffling control showed no such relationships despite preserved label proportions, with both on- and off-Buscar scores remaining uniformly low irrespective of phenotype prevalence (**Supplementary Figure 4**). We next measured the relationship between rankings (sorted by on-Buscar scores) and the proportion of held-out cells labeled for each perturbed gene. Across all 16 phenotypes, gene knockdowns with a higher proportion of cells labeled with a given phenotype achieved significantly lower on-Buscar ranks than genes without such cells (Spearman ρ = −0.52, goodness of fit R^2^ = 0.14, p < 0.0001; **Figure 3B**), confirming that as the fraction of cells per gene displaying a phenotype increases, the gene achieves a lower on-Buscar score and therefore ranks higher in the perturbation ranking for that phenotype. This trend remained consistent across phenotypes, with 13/16 phenotypes (81%) displaying a negative association (**Supplementary Figure 5A**). In contrast, the feature shuffling control removed this relationship (4/16 of phenotypes; 25%) despite preserved label proportions, with both on- and off-Buscar scores remaining uniformly low and the correlation between rank and cell proportion decoupled (Spearman ρ = 0.13, goodness-of-fit R^2^ = 0.00, p = 1.22×10^-4^; **Supplementary Figure 5B**, **Supplementary Figure 6**), confirming that the proportion of labeled cells does not explain gene rank in the absence of morphological signal.

**Figure 3.**
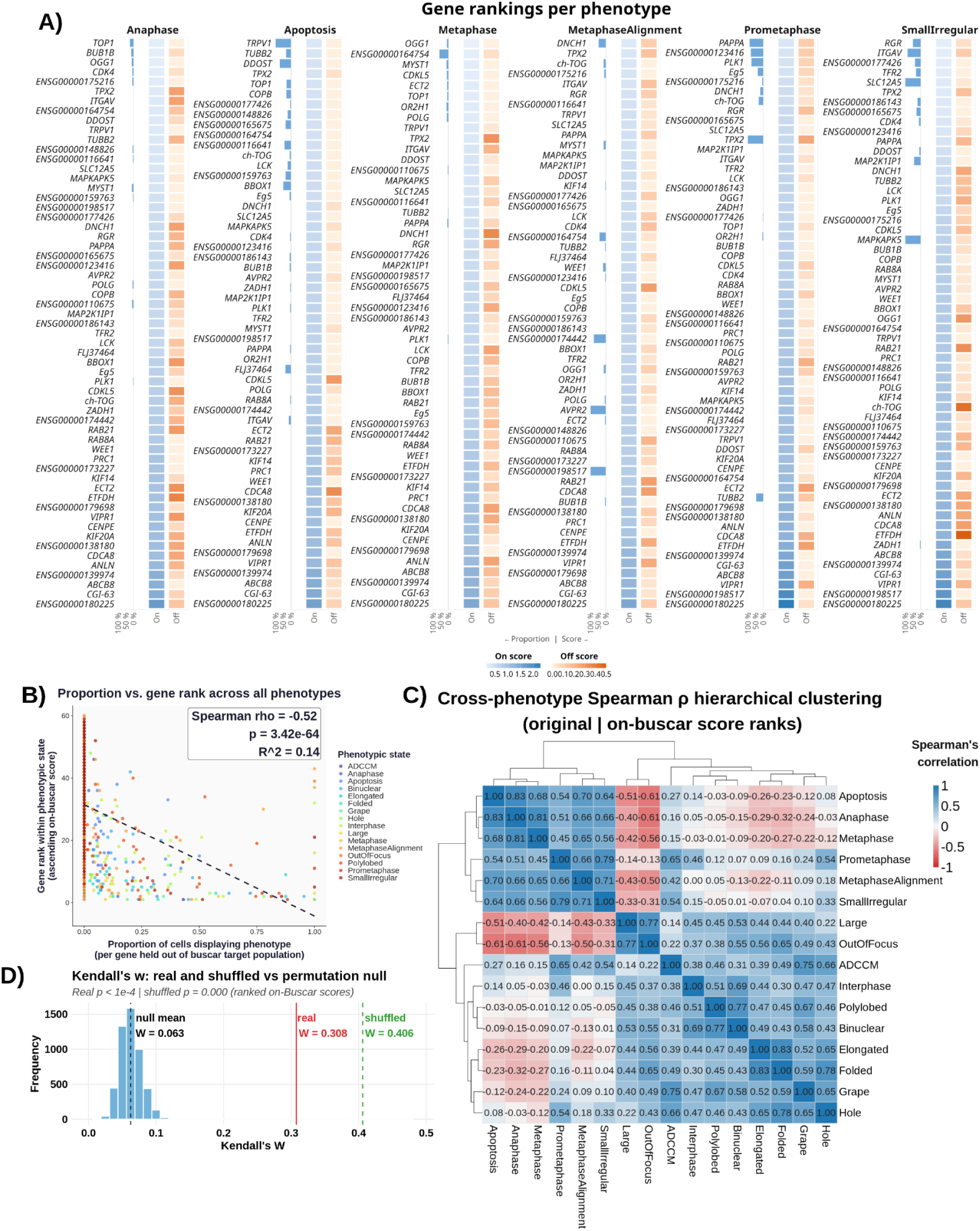
Leave-one-gene-out (LOGO) results for the MitoCheck dataset. **(A)** Heatmaps faceted by Mitocheck’s annotated phenotypes associated with nuclear condensation. Each panel includes a bar plot (left) showing the proportion of cells exhibiting the phenotype of interest for each perturbed gene, alongside two-column heatmaps displaying on-Buscar (blue) and off-Buscar (orange) scores, where darker intensities indicate higher values. We calculated these Buscar scores per phenotype and per gene when all cells of that gene were held out of the target population. We used interphase cells as the reference population. We observed lower on- and off-Buscar scores for genes that contain a higher proportion of single cells labeled with that phenotype. **(B)** Perturbed genes with cells of a given labeled phenotype typically have lower on-Buscar ranks than perturbed genes without cells labeled with that phenotype. **(C)** Each square represents the pairwise correlation between perturbed gene rank vectors (rankings derived from the on-Buscar scores) of two phenotypes. Hierarchical clustering by Ward method. **(D)** Null distribution of Kendall’s W derived from 5,000 label permutations (black), with the mean indicated by a dashed vertical line. The observed Kendall’s W from the original data is shown in red, and the value obtained from feature-wise (column-wise) permutation is shown in green.The separation between the observed W, label permutation null, and feature-shuffled W confirms that Buscar recovers phenotype-specific gene rankings driven by genuine morphological covariance rather than label proportions alone.

To assess the consistency of Buscar’s scores across phenotypes, we computed cross-phenotype Spearman correlations using gene ranks (ranked using on-Buscar scores [**Figure 3C**] and off-Buscar scores, separately**Supplementary Figure 7]**). We assessed overall concordance using Kendall’s W, where a low value reflects genuine phenotype-specific discrimination rather than uniform rankings across phenotypes. The on-Buscar ranking yielded a Kendall’s W of 0.308, which was significantly greater than the label permutation null (null mean W = 0.063; empirical p < 0.0001; **Figure 3D**), demonstrating that Buscar’s rankings are significantly more concordant across phenotypes than expected by chance. To confirm that this concordance reflects genuine morphological covariance rather than label proportions, the feature shuffling control produced a higher Kendall’s W of 0.507 (**Figure 3D**), indicating that destroying the covariance structure generates spurious cross-phenotype concordance. The lower Kendall’s W in the real on-Buscar data relative to the feature-shuffled condition therefore reflects genuine phenotype-specific separation, where perturbed genes are ranked differently across phenotypes in a manner consistent with their distinct morphological roles.

Hierarchical clustering of the on-Buscar cross-phenotype correlation matrix revealed biologically-interpretable structure: anaphase, metaphase, prometaphase, and apoptosis formed a tight positive cluster, likely reflecting their shared morphological phenotype of DNA condensation (Neumann *et al*., 2010; Pau *et al*., 2013; Frolova *et al*., 2023). A second broader grouping encompassed interphase, polylobed, binuclear, ADCCM, elongated, folded, grape, and hole phenotypes, which co-clustered with low-to-moderate positive correlations across pairs. Within this grouping, polylobed and binuclear showed the strongest pairwise correlation, consistent with their shared origin in cytokinesis failure or aberrant nuclear division (Funk *et al*., 2022), while elongated, folded, grape, and hole broadly associated with interphase-like morphological states characterized by relaxed chromatin and expanded cytoplasmic area (Pau *et al*., 2013). However, this broader cluster did not resolve into fully distinct subgroups, likely reflecting the overlapping nuclear morphologies shared across these phenotypes within the nuclear-centric feature space. Notably, large and OutOfFocus exhibited negative correlations with the DNA condensation cluster, likely reflecting an artifact whereby out-of-focus cells appear optically enlarged and dimmer, producing diffuse nuclear profiles that resemble the expanded nuclear morphology of large interphase cells.

The off-Buscar score rankings yielded a Kendall’s W of 0.344, significantly above the label permutation null (null mean W = 0.062; empirical p < 0.0001; **Supplementary Figure 7**). However, the clustering structure observed in the off-Buscar score ranks correlation matrix does not reflect biological phenotype organization. The correlations are uniformly low-to-moderate and positive across all phenotype pairs, with no groupings corresponding to known cell biology. To confirm that this concordance depends on genuine covariance structure, the feature shuffling control produced a substantially higher Kendall’s W of 0.851 (**Supplementary Figure 8**), as destroying the covariance structure underlying the off-morphology signatures caused all perturbations to appear equally disruptive across all phenotypes (**Supplementary Figure 9**), collapsing gene rankings toward a single uniform order and driving the shuffled W toward 1. The lower feature-shuffled W in the on-Buscar condition (0.507) relative to the off-Buscar feature-shuffled condition (0.851) may partly reflect the nuclear-centric feature space used here, where several phenotypic labels share overlapping nuclear morphologies, limiting the degree to which shuffling fully collapses rankings toward uniformity.

### Buscar produces consistent cross-plate rankings in CPJUMP1

To evaluate how Buscar performs when treatment replicates appear across different experimental plates, we applied Buscar to the CPJUMP1 dataset (Chandrasekaran *et al*., 2024). Cross-plate reproducibility is a critical property for any hit calling method because in practice, high-content screens are often run across multiple plates, and a method that produces inconsistent rankings for the same perturbation measured under independent experimental conditions would not be reliable for hit calling. CPJUMP1 is well suited to test this property because it contains single-cell image-based profiles from two cell lines (U2OS and A549) profiled under two perturbation modalities, small-molecule compounds and CRISPR-Cas9 gene knockdowns, where identical treatments appear across multiple plates, enabling a direct assessment of whether Buscar produces consistent on- and off-Buscar scores for the same perturbation measured independently.

To score perturbations across plates, we designated one plate as the reference plate. From this plate, DMSO-treated cells served as the target population and perturbed cells served as the reference population to generate the on- and off-morphology signatures. These signatures were then used by Buscar to score all treatments on the remaining non-reference plates, with on- and off-Buscar scores normalized by the reference distance computed from the reference plate. To establish a negative control, we compared the results across two conditions: paired replicates, where identical perturbations are being measured across plates, and non-paired replicates, where treatment identities on non-reference plates were randomly permuted prior to scoring. This permutation strategy established a null baseline reflecting score distributions expected when treatment identity carries no predictive value across plates. We repeated this procedure until every plate had served as the reference, enabling a comprehensive assessment of Buscar’s replicate stability across all plate combinations.

Across both cell lines and perturbation modalities, paired treatments yielded significantly lower on-Buscar score distributions compared to the randomly permuted non-paired baseline **(Figure 4)**. Specifically, for compound treatments in U2OS cells, the paired on-Buscar score distribution was more tightly concentrated near 1.0 across replicates (SD = 0.175), whereas non-paired scores were inflated above 1.0 and showed a greater spread (SD = 0.656; Mann-Whitney U, p = 8.83×10^-85^, BH-adjusted) **(Figure 4)**. We observed a similar pattern in A549 cells (paired SD = 0.152 vs. non-paired SD = 0.231; p = 7.65×10^-95^, BH-adjusted), confirming that Buscar reliably recovers cross-plate replicate concordance in the compound setting.

**Figure 4.**
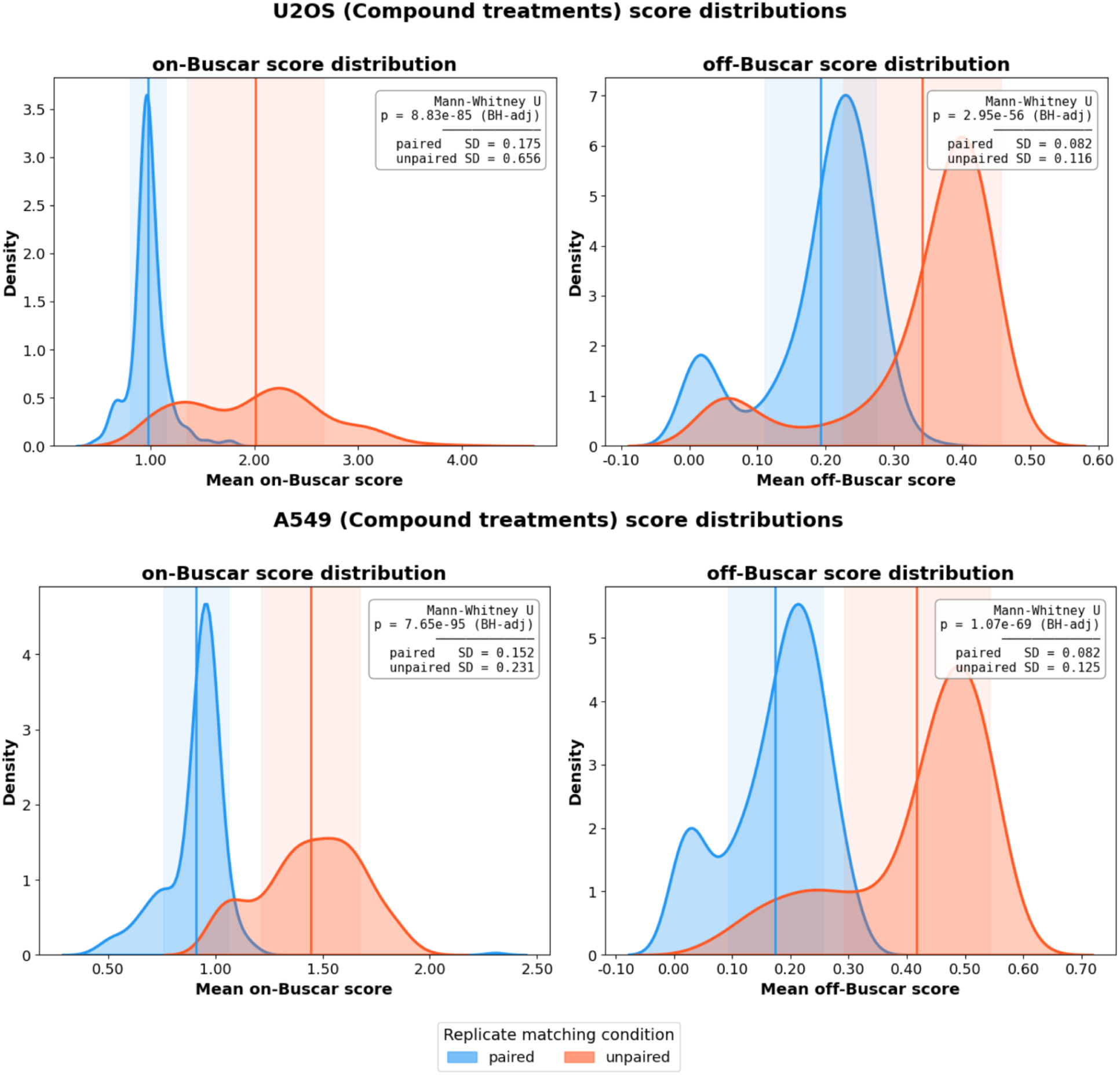
Buscar replicate analysis score distributions for compound perturbations across U2OS and A549 cell lines. Each panel displays kernel density estimates of mean on-Buscar scores (left) and off-Buscar scores (right) per treatment, computed across ten iterations in which every plate serves once as the reference. Distributions are colored by replicate matching condition: blue represents paired perturbation replicates, while orange represents non-paired perturbations (randomly selected perturbations). Vertical lines indicate the mean of each distribution, and shaded regions denote ±1 standard deviation. We assess statistical significance between paired and unpaired distributions using the Mann-Whitney U test with Benjamini-Hochberg (BH) multiple testing correction. Results for small-molecule compound treatments in U2OS (top) and A549 (bottom) cells.

For CRISPR-Cas9 perturbations, we observed the same separation between paired and non-paired on-Buscar score distributions, albeit with a smaller difference in variance relative to compounds **(Supplementary Figure 10)**. This reduced separation likely reflects both the subtler morphology effects typically induced by genetic knockdowns and the smaller number of perturbations profiled (162 gene knockdowns vs. 303 small molecules), which limits the dynamic range of non-paired score distributions. In U2OS cells, paired on-Buscar scores were shifted toward lower values relative to non-paired controls (SD = 0.077 vs. 0.051; p = 1.00×10^-41^, BH-adjusted), and a similar pattern was observed in A549 cells (SD = 0.073 vs. 0.058; p = 2.15×10^-27^, BH-adjusted), indicating that replicate concordance in CRISPR perturbations is reflected primarily in the mean on-Buscar score rather than its variance (**Figure 4**).

Off-Buscar score distributions further support the specificity of Buscar’s cross-plate consistency. Across all conditions, paired treatments consistently produced lower and less dispersed off-Buscar scores relative to non-paired controls. If off-Buscar scores were driven purely by plate-level technical noise rather than perturbation identity, randomly permuting treatment assignments would have no effect on the resulting score distributions, because all treatments would be equally and interchangeably affected by that noise regardless of their identity. The consistent separation observed here instead demonstrates that the off-Buscar score is sensitive to perturbation identity across plates, indicating that off-target morphologies is a reproducible property of the perturbation itself rather than an artifact of the plate on which it was measured. For compound treatments, off-Buscar scores were significantly separated between paired and non-paired conditions in both U2OS (p = 2.95×10^-56^, BH-adjusted) and A549 cells (p = 1.07×10^-69^, BH-adjusted). The same pattern held for CRISPR-Cas9 perturbations, where off-Buscar scores remained significantly separated in U2OS (p = 7.29×10^-28^, BH-adjusted) and A549 cells (p = 1.04×10^-14^, BH-adjusted) (**Figure 4**). Together, these results demonstrate that Buscar robustly recovers cross-plate consistency across both perturbation modalities and cell lines, with compound treatments exhibiting a particularly strong signal consistent with their larger morphological effect sizes.

## Discussion

We present Buscar, a computational method for hit calling in high-content imaging screens that operates directly on distributions of single-cell image-based profiles rather than relying on aggregated summary statistics. By preserving cell-to-cell variability, Buscar addresses a central limitation of conventional image-based profiling approaches, which aggregates heterogeneous cellular responses into single representations and thereby obscures subpopulation structure. Across multiple datasets and perturbation modalities, we demonstrate that leveraging single-cell distributions enables sensitive and interpretable characterization of phenotypic responses, particularly in contexts where treatment effects are subtle, heterogeneous, or context-dependent.

At its core, Buscar simultaneously quantifies perturbation efficacy and specificity through a data-driven separation of the morphology feature space into two complementary signatures derived from two control populations representing distinct phenotypic states. This separation enables the independent tracking of desired morphology shifts through the on-Buscar score and unintended morphology deviations through the off-Buscar score, providing more information about how a perturbation behaves in morphology space than traditional similarity-based hit-calling approaches (Kalinin *et al*., 2025). We recommend prioritizing perturbations with low on-Buscar scores, as these reflect the strongest shift toward the desired phenotypic state. Among perturbations with high efficacy, the off-Buscar score should then be used to deprioritize candidates that broadly alter off-morphology signature features, as perturbations inducing such changes are likely to carry off-target morphological effects that reduce their specificity. Together, these complementary scores allow users to explicitly navigate the trade-off between efficacy and specificity, selecting perturbations that achieve strong phenotypic rescue while minimizing unintended morphological activity for follow-up experimental investigation.

Our benchmarking analyses highlight several desirable properties of Buscar. In the Cell Painting cardiac fibrosis dataset, Buscar captures TGFβRi induced phenotypic rescue in failing cardiac fibroblasts while retaining sensitivity to distinct healthy CF subpopulations and off-target morphological effects specific to the failing cell state that would be obscured by aggregation-based analyses. In the MitoCheck dataset, LOGO evaluation shows that Buscar recovers gene-knockdown phenotype associations in which genes producing a higher proportion of cells with a given nuclear morphology phenotype are consistently ranked above genes that do not, with cross-phenotype rankings reflecting known relationships between mitotic, apoptotic, and nuclear division phenotypes. In the CPJUMP1 dataset, Buscar produces consistent scores for identical perturbations measured across independent plates, including for both small-molecule and CRISPR-based interventions.

Despite these advantages, several limitations remain. First, the current signature construction treats features independently using per-feature statistical testing, which does not account for correlations between features and may therefore limit sensitivity to phenotypes present in multivariate feature interactions. Replacing this per-feature approach with a multivariate two-sample test such as Maximum Mean Discrepancy (MMD) (Gretton *et al*., 2012), which operates on full single-cell distributions without assuming a parametric form would increase sensitivity to complex phenotypes. Second, while the on-Buscar score quantifies the distributional distance between perturbed and target populations, it does not directly measure the proportion of cells that have shifted into the target distributional space. Adding a distribution overlap metric such as the Overlap Coefficient (OVL) (Komaba *et al*., 2023) would complement the on-Buscar score by providing an interpretable estimate of phenotypic rescue, directly quantifying how many cells have returned to the target morphological state. Third, the reliance on EMD introduces computational challenges, as its complexity scales quadratically with the number of cells, which may limit scalability for very large datasets. When using Buscar, we recommend subsampling single cells where possible to reduce computational burden. Future implementations would include approximation methods such as Diffusion EMD (Tong *et al*., 2021) represent a promising direction to fully address this bottleneck without sacrificing distributional sensitivity. Fourth, the quality of the on- and off-morphology signatures depends on the definition and separability of the reference and target populations; poorly defined or highly heterogeneous phenotypes may reduce the discriminative power of Buscar. Finally, while Buscar preserves single-cell heterogeneity, it does not explicitly identify or annotate subpopulations contributing to the observed phenotypic shifts. Explicitly identifying these groups would enable users to directly assess which cell subsets are successfully responding to a perturbation, providing mechanistic insights into drug response heterogeneity (Kinnunen *et al*., 2024).

In summary, Buscar provides a general and extensible method for leveraging single-cell morphology distributions in high-content screening. By jointly quantifying efficacy and specificity through interpretable, distribution-aware metrics, the method offers a principled alternative to aggregation-based approaches and supports accurate and biologically informative hit identification across diverse perturbation types and experimental contexts.

## Methods

### Bioactive Unbiased Single-cell Compound Assessment and Ranking (Buscar)

#### Module 1: Generating on- and off-morphology signatures

Buscar derives on- and off-morphology signatures by comparing single-cell feature distributions between a reference population (e.g., disease cells) and a target population (e.g., healthy cells). For each morphology feature, Buscar applies a non-parametric, two-sample Kolmogorov–Smirnov (KS) test to assess whether the feature distribution differed significantly between the two populations. The KS statistic quantifies the maximum absolute difference between the empirical cumulative feature distribution of the reference cell population against that of the target population. To control for multiple comparisons across the full feature space, Buscar corrects p-values for multiple tests using the Benjamini-Hochberg false discovery rate (FDR) procedure at a significance threshold α = 0.05.

Buscar uses the KS-test results to define the signatures. Specifically, Buscar adds morphology features with FDR-corrected p-values below this threshold, indicating statistically significant distributional shifts between the reference and target populations, to the on-morphology signature. The on-morphology signature represents morphologies altered in the target condition of interest. Typically, the target condition will represent a healthy phenotype, and therefore, the significantly different morphologies represent which features must change in order for a perturbation (typically in a high-throughput experiment) to revert the disease condition to a healthy state. Conversely, Buscar assigns features with corrected p-values at or above the threshold, indicating no statistically-detectable difference between populations, to the off-morphology signature. The off-morphology signature represents morphologies unrelated to the target phenotype. In other words, if a perturbation alters these features, it would indicate off-target effects of the perturbation. In summary, this process yields two complementary feature sets: on-morphology features used to quantify phenotypic reversal, and off-morphology features for detecting off-target effects.

#### Module 2: Perturbation efficacy and specificity scoring

Buscar quantifies perturbation efficacy (on-Buscar score) and specificity (off-Buscar score) in a two-step procedure. Step 1: Buscar calculates a reference distance between the reference state and a target state using the on-morphology feature set. Buscar uses this distance to normalize all perturbation scores where a score close to 1.0 indicates that cells remain morphologically similar to the reference state, reflecting low efficacy, while a score close to 0.0 indicates that cells have shifted toward the target state, reflecting high efficacy. Scores greater than 1.0 are also possible and indicate that the perturbation has pushed cells further away from the target state than the reference cells themselves.

Step 2: Buscar computes two scores for each perturbation, the on-Buscar score and the off-Buscar score. Buscar computes the on-Buscar score using the Earth Mover’s Distance (EMD), also known as the Wasserstein distance, applied exclusively to the on-morphology feature set. EMD quantifies the minimum amount of work required to transform the single-cell feature distribution of the perturbed population into the target population (Orlova *et al*., 2016). In practice, due to intrinsic cell heterogeneity, a score of exactly 0.0 is not strictly attainable and observed scores approach but do not reach zero.

Buscar next computes the off-Buscar score by applying a KS test to each feature in the off-morphology signature, comparing the reference and perturbed populations. Features that become statistically significant after FDR correction are interpreted as features unintentionally altered by the perturbation. The off-Buscar score is the proportion off-morphology features that become significant over the total number of features in the off-morphology signature, where a low score indicates that the perturbation does not broadly alter off-target cell morphology features.

#### Compound scoring in the Cell Painting cardiac fibroblasts dataset

The dataset comprises image-based profiles derived from two biological conditions: Cardiac fibroblasts (CFs) extracted from healthy human hearts and hearts from patients with cardiac fibrosis. Each heart was treated with either a vehicle control (DMSO) or transforming growth factor beta receptor inhibitor (TGFβRi). This experimental design yields four distinct groups: (i) healthy CFs treated with DMSO, (ii) healthy CFs treated with TGFβRi, (iii) failing CFs treated with DMSO, and (iv) failing CFs treated with TGFβRi. The dataset contains a total of 15,793 single-cell profiles distributed across six replicate wells for healthy DMSO, failing DMSO, and healthy TGFβRi conditions, and 15 replicate wells for the failing TGFβRi condition. For full details about this dataset see (Travers *et al*., 2025).

We applied Buscar to this dataset as a pilot experiment. First, we applied Buscar’s Module 1 (generating on- and off-morphology signatures), defining the reference population as DMSO-treated failing CFs and the target state as DMSO-treated healthy CFs. The failing CF reference population was derived from patients diagnosed with idiopathic dilated cardiomyopathy (Travers *et al*., 2025). We next applied Buscar Module 2 to compute on-Buscar scores (efficacy) and off-Buscar scores (specificity) for all four perturbations: healthy DMSO (the target state), failing DMSO (the reference state), and healthy TGFβRi and failing TGFβRi (both representing perturbed states).

We generated UMAP embeddings separately for the on- and off-morphology signatures to visualize how each signature captures variation in single-cell phenotypes. Specifically, we performed dimensionality reduction on single-cell profiles using only the features corresponding to each signature subset. For both embeddings, we computed UMAP using two components (n_components = 2) using the cosine distance metric and a fixed random seed to ensure reproducibility. We left all other parameters at their default settings as implemented in the Python umap-learn package.

#### Mitocheck Leave-One-Gene-Out Analysis

The MitoCheck dataset originates from a genome-wide phenotypic screen in which each of the approximately 21,000 human protein-coding genes was individually silenced by RNA interference (RNAi) in HeLa cells stably expressing histone H2B fused to green fluorescent protein, followed by two-day time-lapse live imaging of fluorescently labelled chromosomes (Neumann *et al*., 2010). The full dataset is publicly available through the Image Data Resource (IDR) under accession idr0013 (screenA) (Williams *et al*., 2017). We used the labelled subset of this dataset, comprising cells with confidently manually-annotated nuclear morphology phenotypes. This labelled subset spans 16 distinct phenotypic states, anaphase, metaphase, prometaphase, apoptosis, binuclear, polylobed, elongated, folded, large, ADCCM, hole, interphase, grape, OutOfFocus, SmallIrregular, and MetaphaseAlignment, each treated as an independent phenotypic category in all downstream analyses. The labelled dataset contains a total of 1,394,555 cells, distributed across three categories: 779,993 negative control cells (scrambled siRNA), 612,059 positive control cells corresponding to knockdown of the mitotic regulators KIF11, COPB1, and ENSG00000149503, and 2,503 phenotypically labeled cells. For full image-based profiling processing details of this dataset see (Tomkinson *et al*., 2024). In this paper, we only analyzed the negative control cells (scrambled siRNA) and the 2,503 manually annotated phenotypically labeled cells.

A key requirement for a hit calling method is the ability to recover biologically meaningful gene-phenotype associations, that is, to rank genes whose knockdown produces a given phenotype more highly than genes whose knockdown does not. To evaluate whether Buscar meets this requirement, we used the MitoCheck dataset, which enables a direct assessment of whether Buscar’s scores reflect known biological signals. A concern in this setting is data leakage: if cells from the same gene knockdown appear in both the target population used to define the morphology signature and the query population being scored, Buscar would trivially recover that gene due to shared identity rather than genuine morphology signal. To prevent this, we performed a leave-one-gene-out (LOGO) analysis, in which all cells from a given gene knockdown were held out from the target population before scoring, ensuring that Buscar’s rankings reflect morphological generalization rather than memorization of training signal

The LOGO procedure consisted of the following steps. For each of the 16 phenotypes independently, we divided perturbed genes into two groups: those whose knockdown produced cells labeled with the given phenotype and those whose knockdown did not. For each phenotype, we applied Buscar once per gene in the first group, holding out all cells from that gene knockdown prior to scoring to prevent data leakage. In all cases, we used interphase cells (from scramble siRNA control) as the reference state and all remaining single cells labeled with that phenotype, excluding cells of the held-out gene, as the target state. We then scored all remaining cells, treating each gene as a distinct perturbation group. We excluded gene knockdowns with fewer than five single-cell profiles in the phenotype of interest due to the target population being too small to produce stable feature distributions required for on-Buscar scoring (which uses EMD), or where no features passed the FDR threshold, resulting in empty on- or off-morphology signatures.

To establish a negative control baseline, we applied the same LOGO analysis to both the original profiles and a feature-shuffled version. For the shuffled condition, we permuted morphology feature values column-wise across all single-cell profiles in order to remove any biological signal from the feature space while leaving gene and phenotype labels unaffected, and randomly reassigned features between the on- and off-morphology signatures while preserving the original size of each signature

To assess whether Buscar produces phenotype-specific gene rankings rather than ranking genes uniformly regardless of phenotypic state, we computed Kendall’s W across all 16 phenotypes for genes with complete rankings, where a low value indicates that gene rankings differ across phenotypes reflecting genuine phenotype-specific discrimination, and a high value indicates that genes are ranked similarly regardless of phenotypic state. To assess statistical significance, we compared the observed Kendall’s W against an empirical null distribution that we generated by randomly permuting gene labels 5,000 times and recalculating Kendall’s W for each permutation, defining the empirical p-value as the proportion of permuted W values greater than or equal to the observed value. We computed pairwise Spearman rank correlations between per-phenotype gene ranking for all phenotype pairs to assess the structure of phenotypic similarity in the ranking space. Finally, to quantify the relationship between the proportion of phenotype-labeled cells per perturbed gene and its on-Buscar score rank, we computed the Spearman correlation between these two variables across all phenotypes.

#### CPJUMP1 replicate analysis

We used the CPJUMP1 dataset (Chandrasekaran et al., 2024), which profiles two perturbation modalities, small-molecule compounds and CRISPR-Cas9 gene knockdowns, across two cell lines (A549 and U2OS) and includes a series of experimental controls distributed across plates: negative controls consisting of DMSO-treated wells for compound perturbations and non-targeting single guide RNA (sgRNA) wells for CRISPR-Cas9 perturbations. We reprocessed the CPJUMP1 dataset using CytoTable, coSMicQC, and Pycytominer, and we made the pipeline available at: https://github.com/WayScience/JUMP-single-cell. We used feature-selected profiles for all downstream analyses. We subset profiles by cell line, selecting plates with matched experimental conditions: seeding density of 100% for both perturbation modalities, incubation times of 48 hours for small-molecule compounds and 144 hours for CRISPR-Cas9 perturbations, and no antibiotic selection. This yielded four plates per cell line per perturbation modality, for a total of 16 plates. For compound treatments, our matched plates contained 303 small molecules shared across both cell lines, while for CRISPR-Cas9 perturbations, our matched plates contained 162 gene knockdowns shared across both cell lines. The final dataset comprised 1,135,692 single-cell profiles for compound-treated U2OS cells, 3,485,667 for compound-treated A549 cells, 1,349,188 for CRISPR-Cas9-treated U2OS cells, and 1,525,556 for CRISPR-Cas9-treated A549 cells.

To evaluate the cross-plate reproducibility of Buscar across perturbation modalities, we conducted a replicate consistency analysis using the CPJUMP1 dataset. For each cross-plate comparison, we designated one plate as the reference plate. From this plate, we derived on-and off-morphology signatures using Buscar’s Module 1 by comparing the perturbed cells (reference) to the control cells (target). To ensure robustness to control sampling, we performed this signature derivation step across 10 independent iterations, where the control cells were randomly subsampled by 0.02% for each iteration. We then used the resulting signatures to score all treatments on the remaining non-reference plates. The resulting on- and off-Buscar scores were normalized by the reference plate distance such that a score of 1.0 corresponds to the morphological distance of the treatment on the reference plate, and a score of 0.0 corresponds to the baseline defined by the negative control. We repeated this procedure until each of the four plates had served as the reference, generating a fully-crossed set of 12 cross-plate comparisons per treatment for each cell line and perturbation modality.

To establish a negative control distribution reflecting the expected score range in the absence of replicate concordance, we randomly permuted treatment identities on non-reference plates prior to scoring for each reference plate and treatment. We repeated this permutation in 10 independent iterations per plate pair, generating a null distribution of on- and off-Buscar scores against which we compared paired replicate scores. We evaluated statistical differences between paired and non-paired score distributions using the Mann-Whitney U test with Benjamini-Hochberg correction for multiple comparisons (Haynes, 2013; Mann and Whitney, 1947).

## Data Availability

The Buscar implementation can be found along with all analysis notebooks used in this study at https://github.com/WayScience/buscar.

## Acknowledgements

Research reported in this publication was supported by the National Library of Medicine of the NIH under award number T15LM009451 to E.S., The Gilbert Family Foundation (923014 to G.P.W.), Alex’s Lemonade Stand Foundation ‘A’ Award and Tap Cancer Out (Grant # 23–28306 to G.P.W.), American Heart Association Collaborative Sciences Award (24CSA1255857 to G.P.W.).

