## Supplemental Material for "Single-cell hit calling in high-content imaging screens with Buscar"

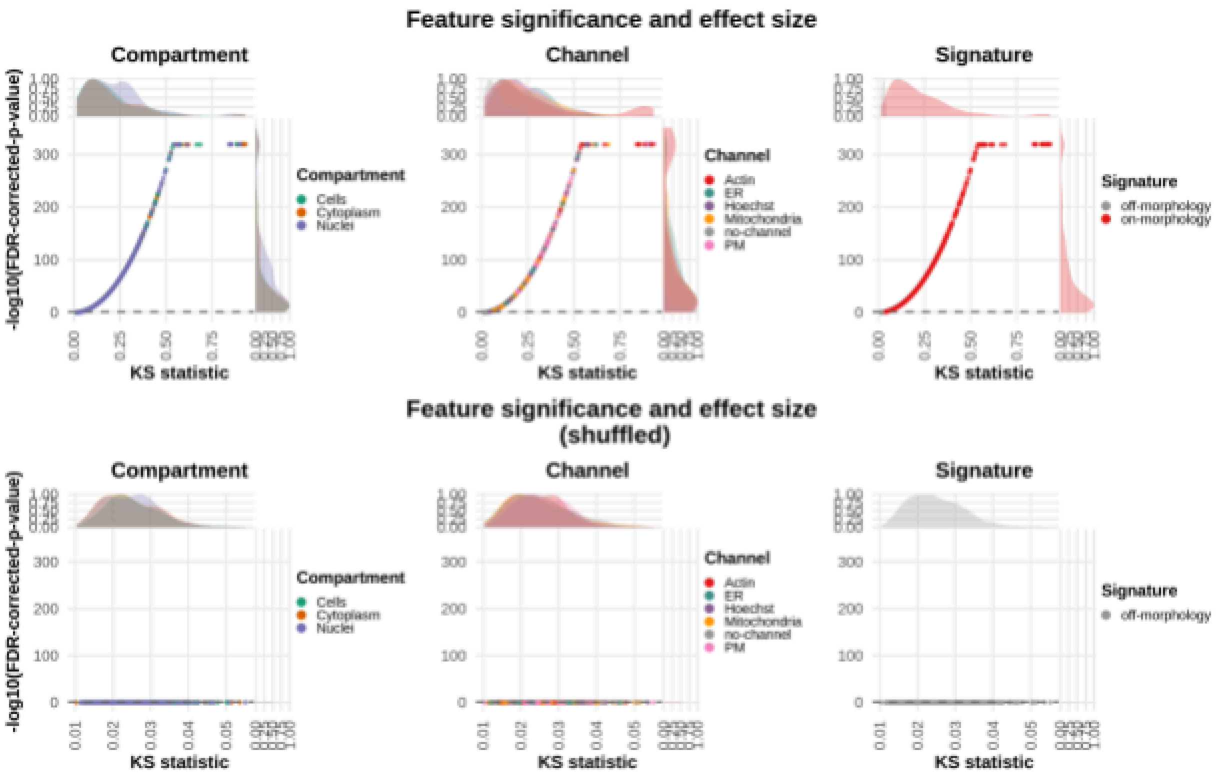

**Supplementary Figure 1. Significance and effect size of on- and off-morphology signatures on shuffled Cell Painting data of healthy and failing cardiac fibroblasts.** We randomly shuffled the healthy and failing CF label and then ran the Buscar signature module. Scatter plots of FDR-corrected p-values (−log10, y-axis) versus Kolmogorov-Smirnov (KS) statistic (x-axis) for each CellProfiler morphology feature, stratified by compartment (nucleus, cytoplasm, and whole cell [left]), channels and by signature type (on- and off-morphology signatures). The marginal density plots (KDE) on the axes represent the relative frequency and concentration of features across different effect size and significance ranges. Buscar correctly assigned all features to the off-morphology score after the shuffling procedure.

| target | perturbation | on_buscar_scores | off_buscar_scores |
|---|---|---|---|
| healthy_DMSO | healthy-TGFRi-C04 | 0.7952824718 | 0.05882352941 |
| healthy_DMSO | failing-TGFRi-B07 | 0.9465550621 | 0.1176470588 |
| healthy_DMSO | failing-TGFRi-D07 | 1.079648315 | 0.2647058824 |
| healthy_DMSO | healthy-TGFRi-E07 | 0.8824083445 | 0 |
| healthy_DMSO | healthy-TGFRi-C07 | 0.7789722459 | 0.05882352941 |
| healthy_DMSO | failing-TGFRi-D10 | 0.9292801373 | 0.1470588235 |
| healthy_DMSO | healthy-TGFRi-E10 | 0.8171432504 | 0.4411764706 |
| healthy_DMSO | failing-TGFRi-B04 | 0.8996053887 | 0 |
| healthy_DMSO | healthy-TGFRi-E04 | 0.9269512768 | 0 |
| healthy_DMSO | healthy-TGFRi-C10 | 0.7713593194 | 0.1176470588 |
| healthy_DMSO | failing_DMSO | 1 | 0 |
| healthy_DMSO | failing-TGFRi-D04 | 0.9478040826 | 0.2352941176 |
| healthy_DMSO | failing-TGFRi-B10 | 0.8742935198 | 0.1176470588 |

**Supplementary Table 1. Pooled Buscar scores for the Cell Painting dataset of cardiac fibroblasts.**
We generated on-Buscar and off-Buscar scores using the reference distance established between the failing CF (reference) cells and healthy CF cells (target) treated with DMSO. All Buscar scores were normalized by this reference distance.

| target | treatment | well | on_buscar_scores | off_buscar_scores | cell_counts |
|---|---|---|---|---|---|
| healthy_DMSO | healthy-TGFRi | C10 | 0.7713593194 | 0.1176470588 | 314 |
| healthy_DMSO | failing-TGFRi | B07 | 0.9465550621 | 0.1176470588 | 545 |
| healthy_DMSO | healthy-TGFRi | C07 | 0.7789722459 | 0.05882352941 | 492 |
| healthy_DMSO | failing-TGFRi | D07 | 1.079648315 | 0.2647058824 | 729 |
| healthy_DMSO | healthy-TGFRi | E07 | 0.8824083445 | 0 | 63 |
| healthy_DMSO | healthy-TGFRi | E10 | 0.8171432504 | 0.4411764706 | 235 |
| healthy_DMSO | failing-TGFRi | D04 | 0.9478040826 | 0.2352941176 | 870 |
| healthy_DMSO | failing-DMSO | | 1 | 0 | 9153 |
| healthy_DMSO | failing-TGFRi | D10 | 0.9292801373 | 0.1470588235 | 685 |
| healthy_DMSO | healthy-TGFRi | C04 | 0.7952824718 | 0.05882352941 | 288 |
| healthy_DMSO | failing-TGFRi | B10 | 0.8742935198 | 0.1176470588 | 618 |
| healthy_DMSO | healthy-TGFRi | E04 | 0.9269512768 | 0 | 94 |
| healthy_DMSO | failing-TGFRi | B04 | 0.8996053887 | 0 | 341 |

**Supplementary Table 2. Replicate-level Buscar scores for the Cell Painting dataset of cardiac fibroblasts.**

Each row represents an individual well replicated score independently. The reference distance was established by pooling all replicates from the failing CF (reference) and healthy CF (target) DMSO-treated populations; this pooled distance was then used to normalize all per-replicate on- and off-Buscar scores.

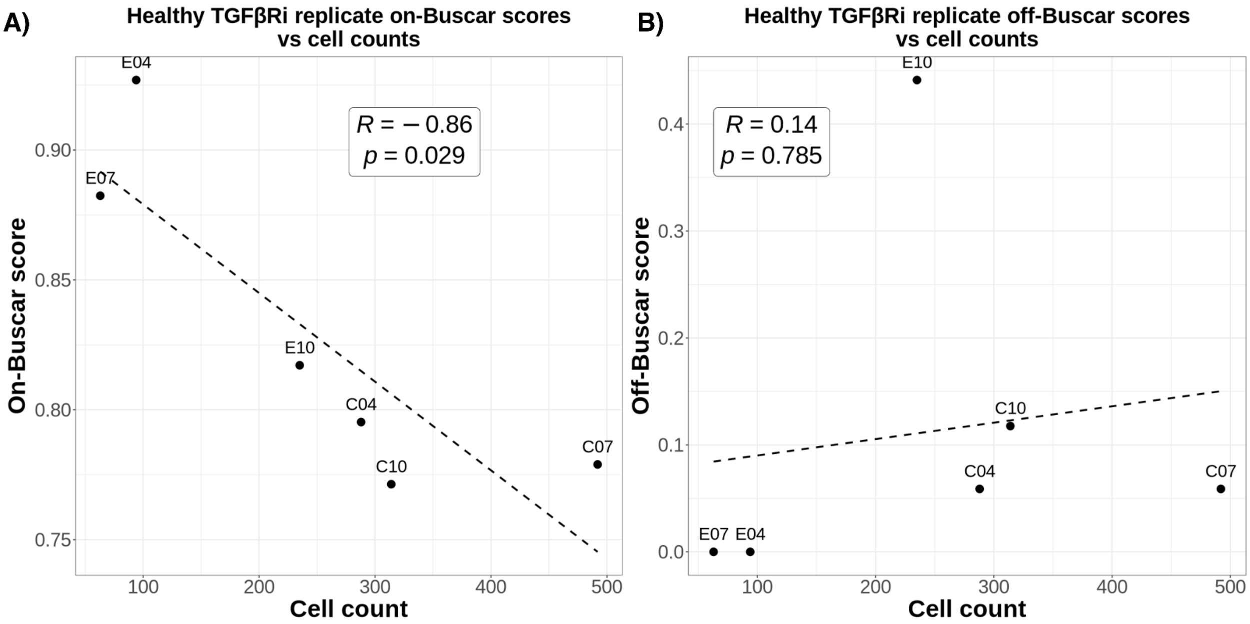

**Supplementary Figure 2. Relationship between well-level cell counts and on and off-Buscar scores for healthy CFs treated with TGFβRi.**
Each point represents an individual replicate well (e.g., Well E07). **(A)** On-Buscar scores for healthy CFs treated with TGFβRi are negatively correlated with cell count (Spearman R = −0.86, p = 0.029). **(B)** Off-Buscar scores show no significant correlation with cell count (Spearman R = 0.14, p = 0.785).

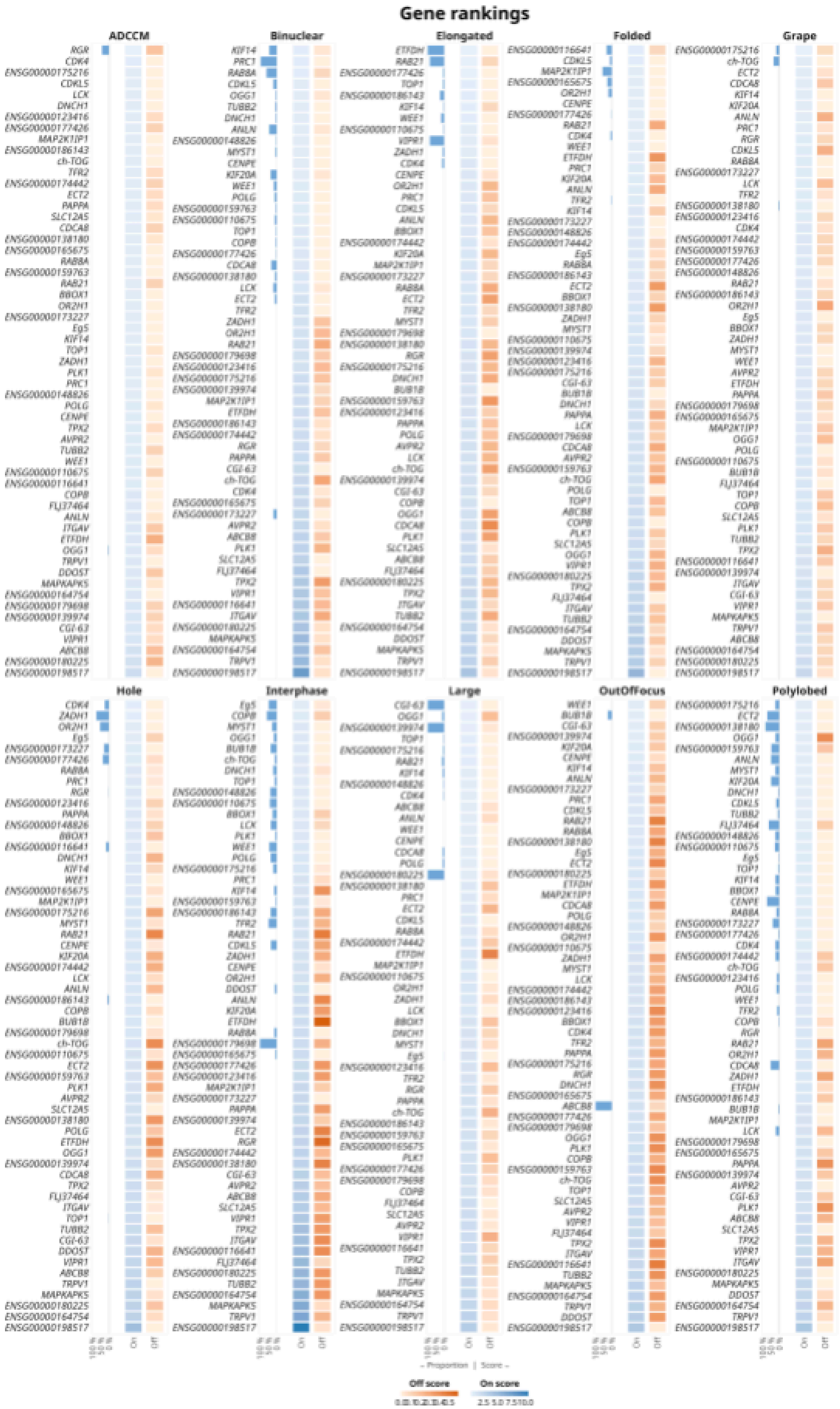
Gene rankings
ADCCM
Binuclear
Elongated
Folded
Grape
Hole
Interphase
Large
OutOfFocus
Polylobed
On
Off
Proportion | Score
Off score
On score

**Supplementary Figure 3. Leave-one-gene-out (LOGO) results for the MitoCheck dataset for interphase-type and polynucleated-type phenotypes.**
Heatmaps faceted by MitoCheck's annotated phenotype for phenotypes associated with interphase (top) and polynucleated (bottom) cell states. Each panel includes a bar plot (left) showing the proportion of cells labeled with the phenotype of interest for each perturbed gene, alongside two-column heatmaps displaying on-Buscar (blue) and off-Buscar (orange) scores, where darker intensities indicate higher values. We calculated these Buscar scores per phenotype and per gene when all cells of that given gene were held out of the target population. We used interphase cells as the reference population. See methods for complete details.

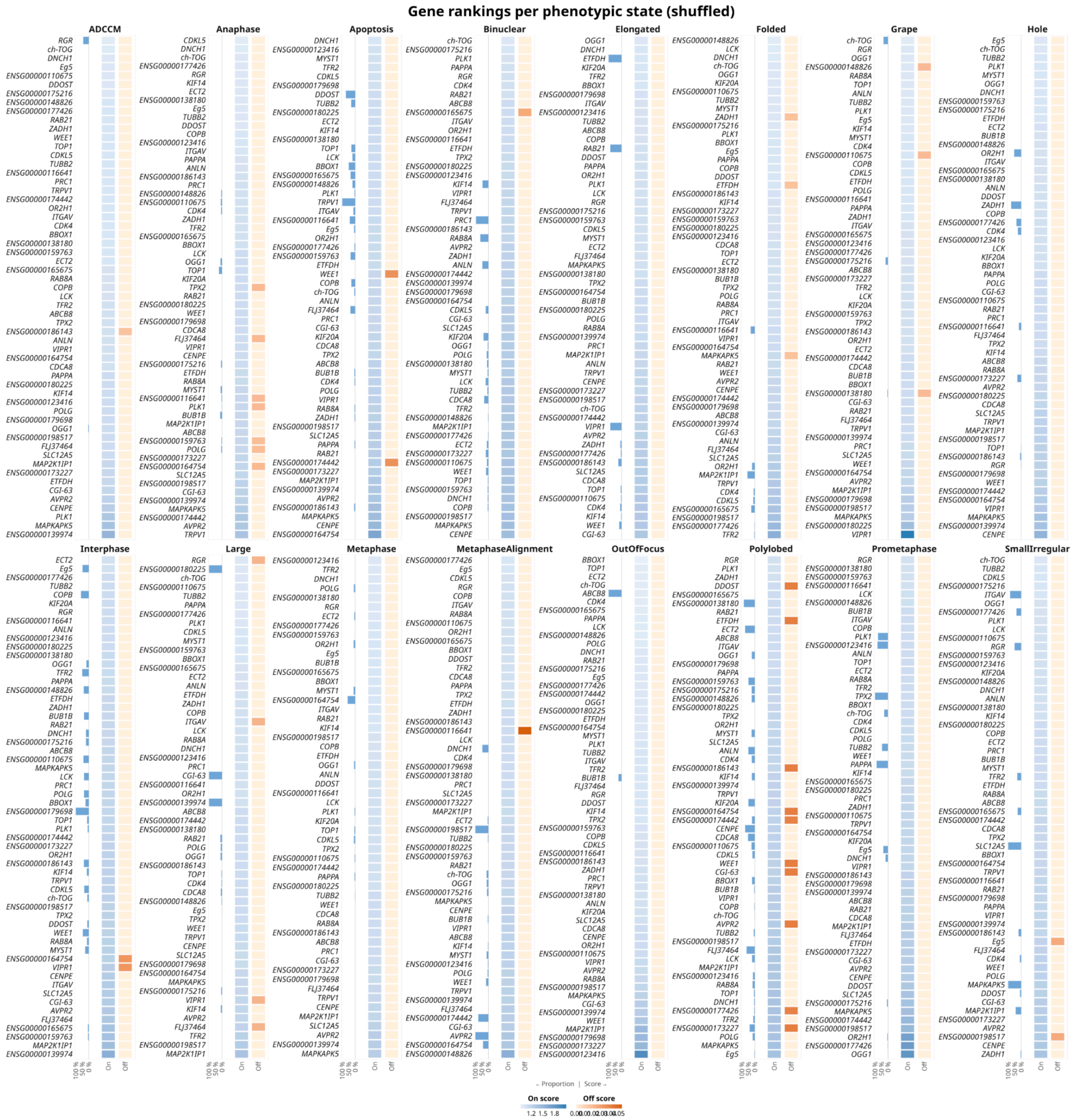

**Supplementary Figure 4. Feature-shuffled leave-one-gene-out (LOGO) results for the MitoCheck dataset.**

Heatmaps faceted by MitoCheck's labeled phenotype. Each panel includes a bar plot (left) showing the proportion of cells exhibiting the phenotype of interest for each perturbed gene, alongside two-column heatmaps displaying on-Buscar (blue) and off-Buscar (orange) scores, where darker intensities indicate higher values. All profiles were feature-shuffled prior to scoring by permuting morphological feature values column-wise, destroying the real covariance structure while preserving gene labels and phenotype proportions. We used interphase cells as the reference population. See methods for complete details.

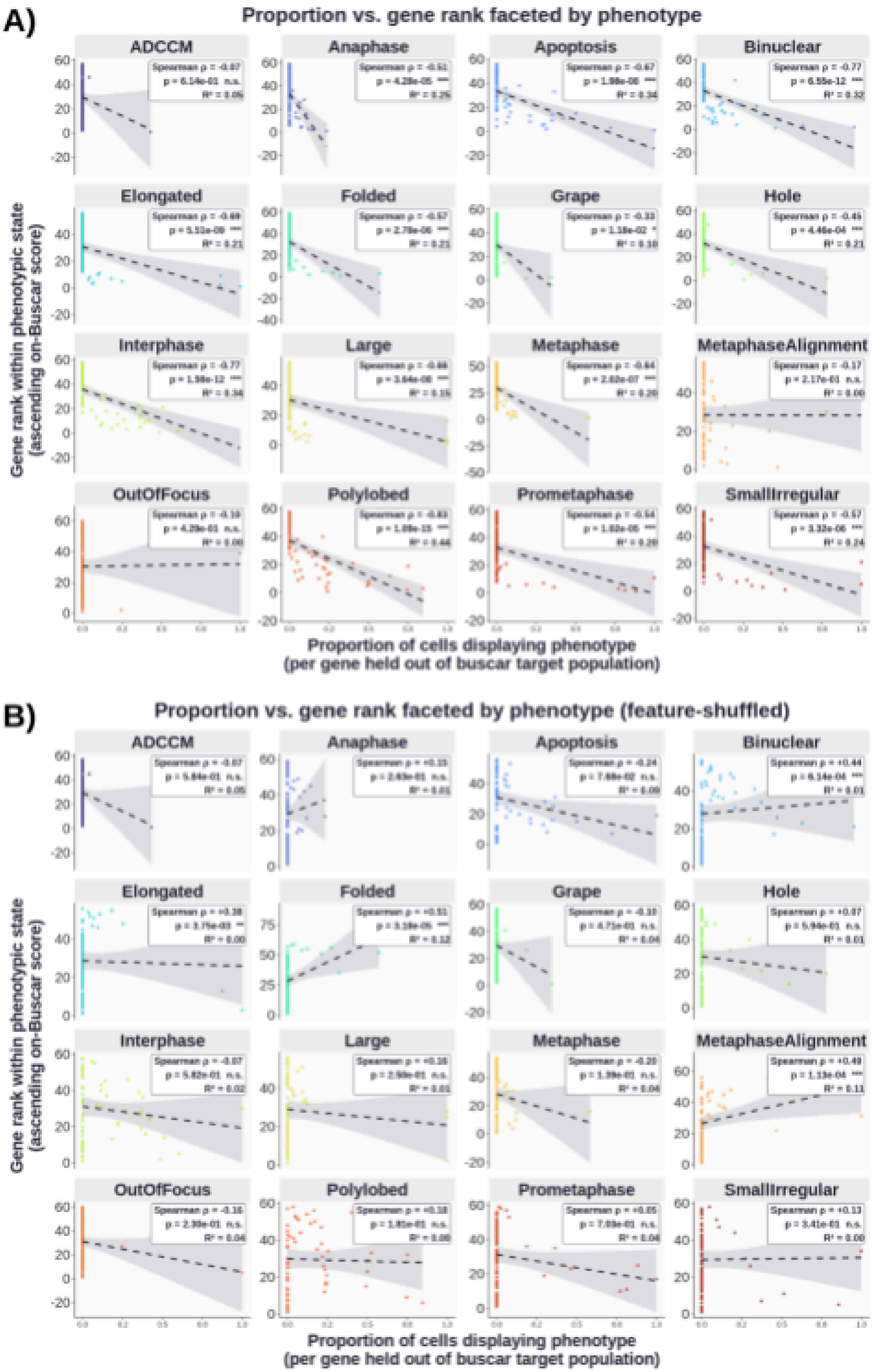

**Supplementary Figure 5. Proportion of phenotype-labeled cells versus Buscar gene rank for real and feature-shuffled data.**

Scatter plots show gene rank (ascending on-Buscar score, y-axis) against the proportion of held-out cells displaying each phenotype (x-axis), faceted by phenotypic state, for **(A)** original and **(B)** feature-shuffled profiles, where we randomly permuted morphology feature values column-wise prior to scoring. Each point represents a perturbed gene; dashed lines indicate linear regression fits. Spearman ρ, Benjamini-Hochberg adjusted p-values, and goodness-of-fit $R^2$ are reported per panel. Significance: n.s. not significant, * $p < 0.05$, ** $p < 0.01$, *** $p < 0.001$.

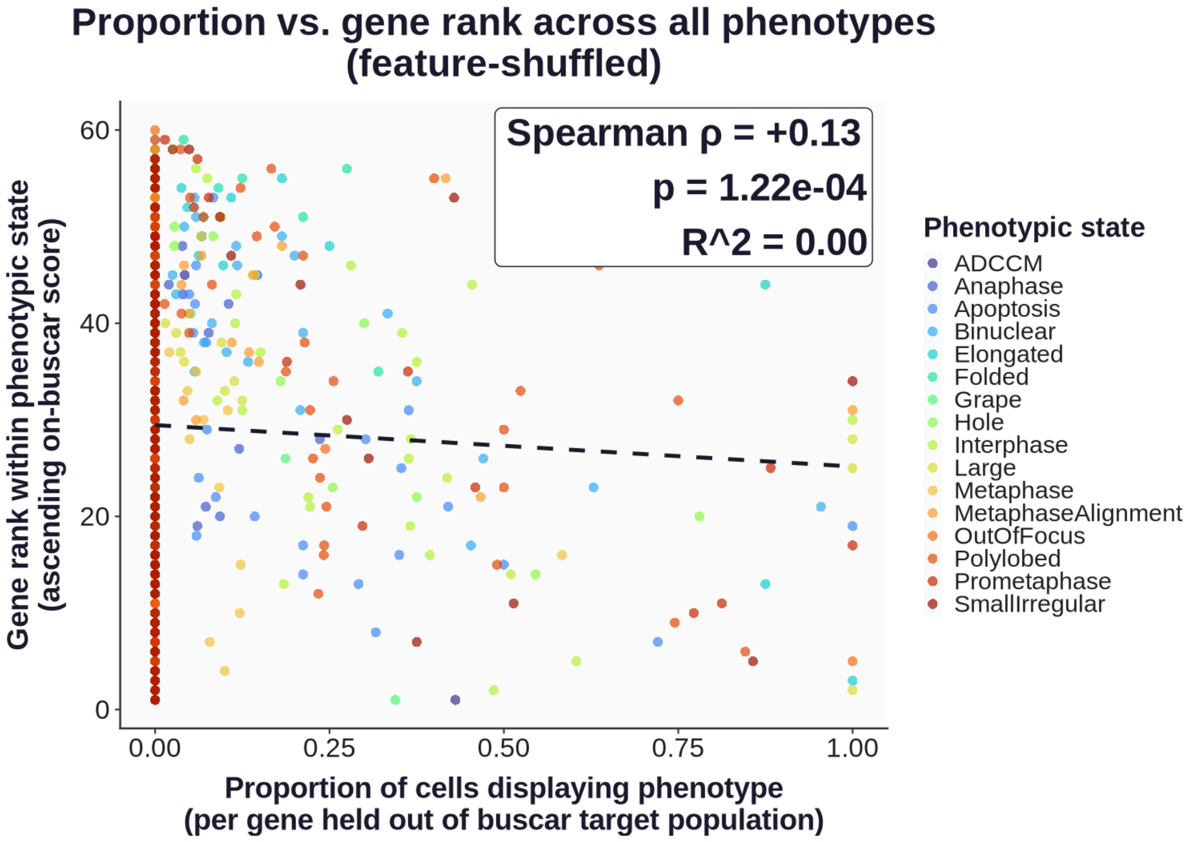

**Supplementary Figure 6. Proportion vs. gene rank across all phenotypes under feature-shuffled conditions.**

Each point represents a perturbed gene within a given phenotypic state, with the proportion of cells displaying the phenotype of interest (per gene held out of the Buscar target population) on the x-axis and the gene rank derived from ascending on-Buscar scores on the y-axis. Points are colored by phenotypic state. The dashed line indicates the linear regression fit. Morphological feature values were permuted column-wise prior to scoring.

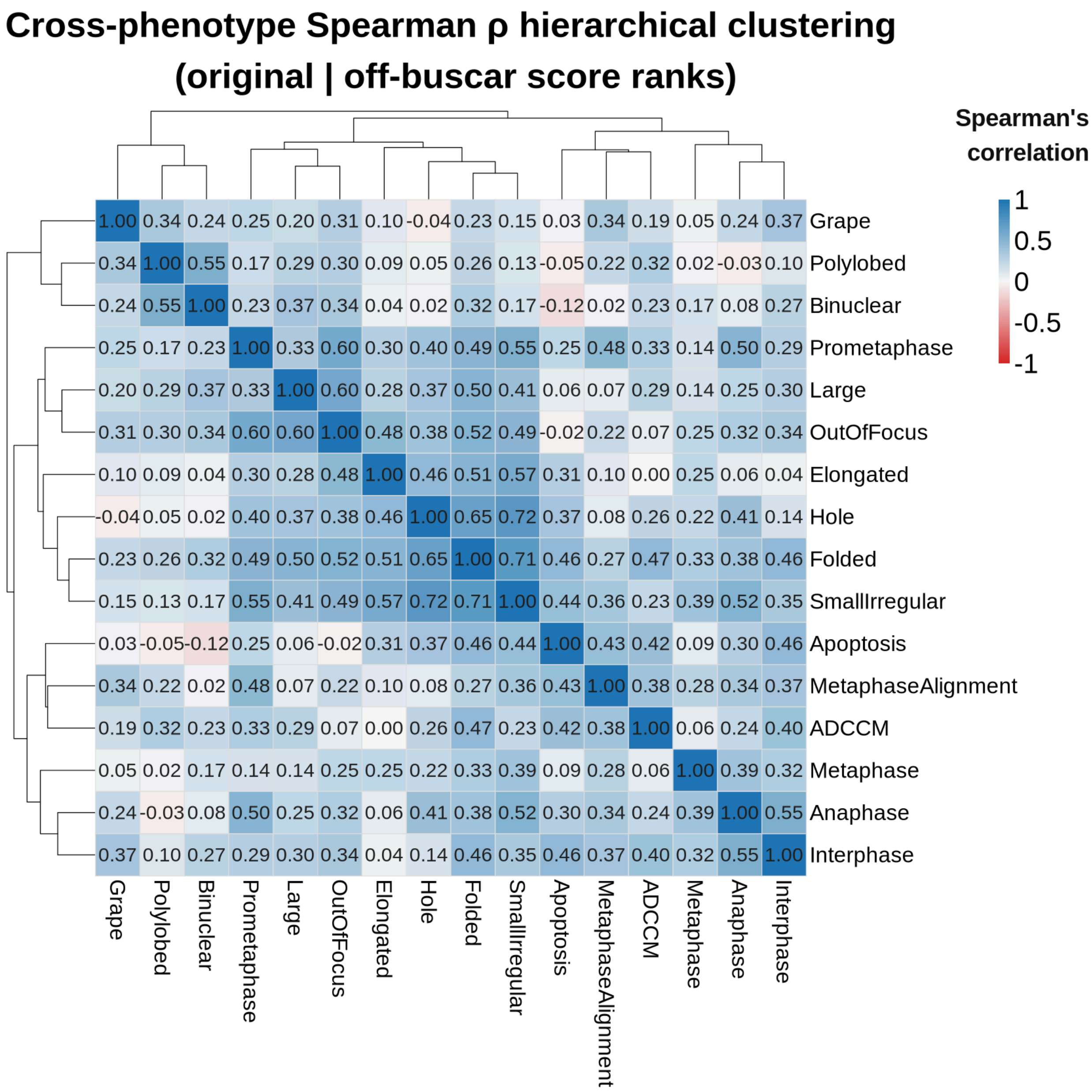

**Supplementary Figure 7. Cross-phenotype Spearman ρ hierarchical clustering based on off-Buscar scores.**

Hierarchically clustered correlation matrix where each cell represents the pairwise Spearman rank correlation between gene ranking vectors of two phenotypes, derived from off-Buscar scores across 16 phenotypes. Color scale indicates Spearman's correlation ranging from −1 (red) to +1 (blue). Hierarchical clustering was performed using the Ward method.

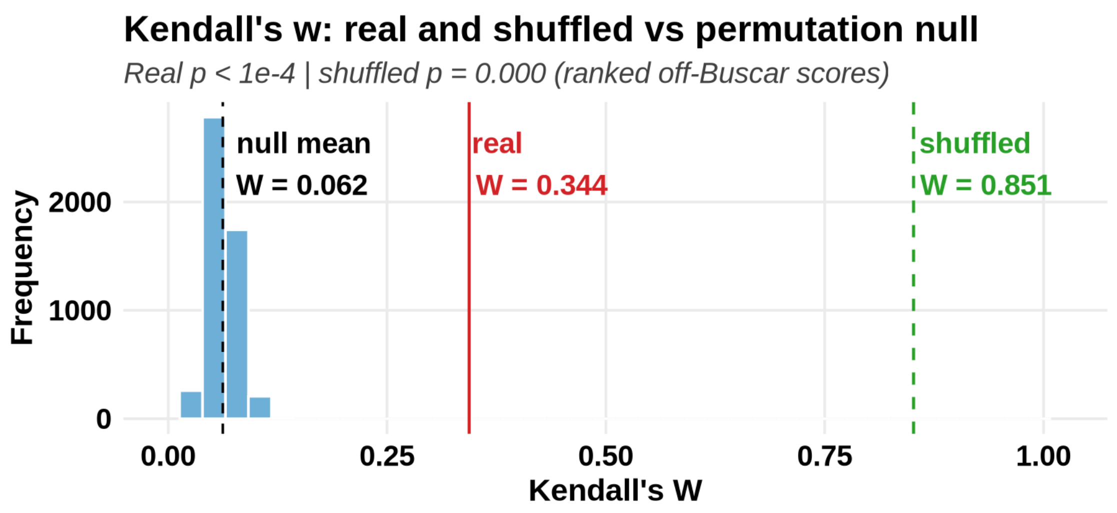

**Supplementary Figure 8. Kendall's W concordance analysis for off-Buscar score rankings.**
Null distribution of Kendall's W derived from 5,000 label permutations (black), with the mean indicated by a dashed vertical line. The observed Kendall's W from the original data is shown in red, and the value obtained from feature-wise (column-wise) permutation is shown in green.

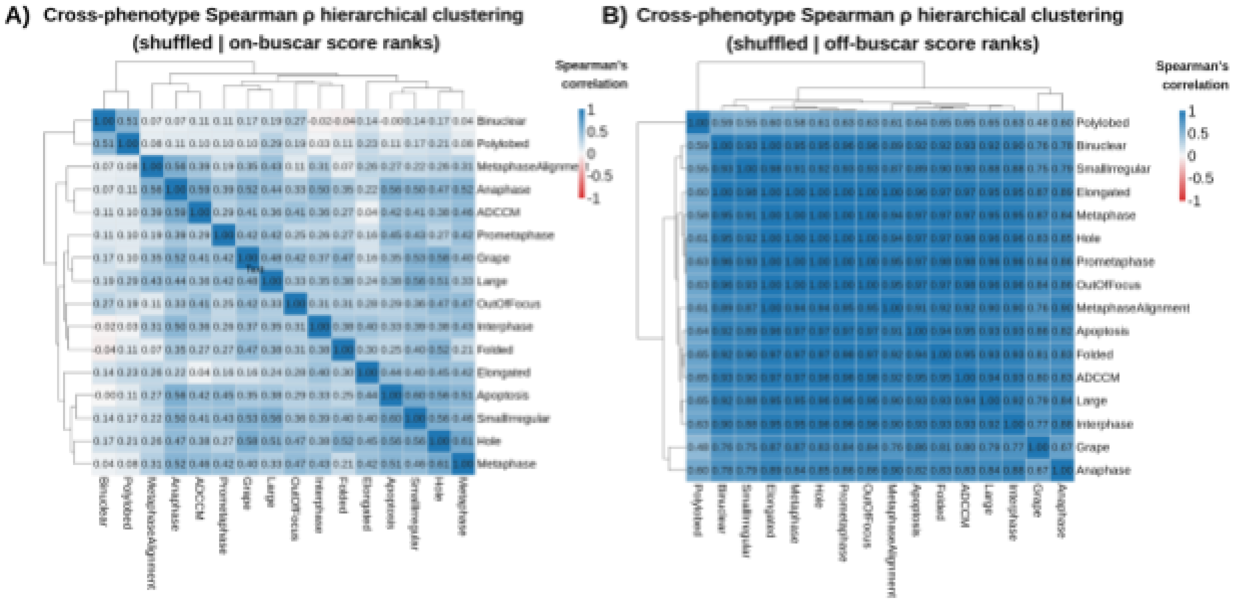

**Supplementary Figure 9. Cross-phenotype Spearman ρ hierarchical clustering under feature-shuffled conditions.**

Each square represents the pairwise Spearman correlation between perturbed gene rank vectors across 16 phenotypes, derived from on-Buscar (A) and off-Buscar (B) score ranks under feature-shuffled conditions, where morphological feature values were permuted column-wise prior to scoring. Color scale indicates Spearman's correlation ranging from −1 (red) to +1 (blue). Hierarchical clustering was performed using the Ward method.

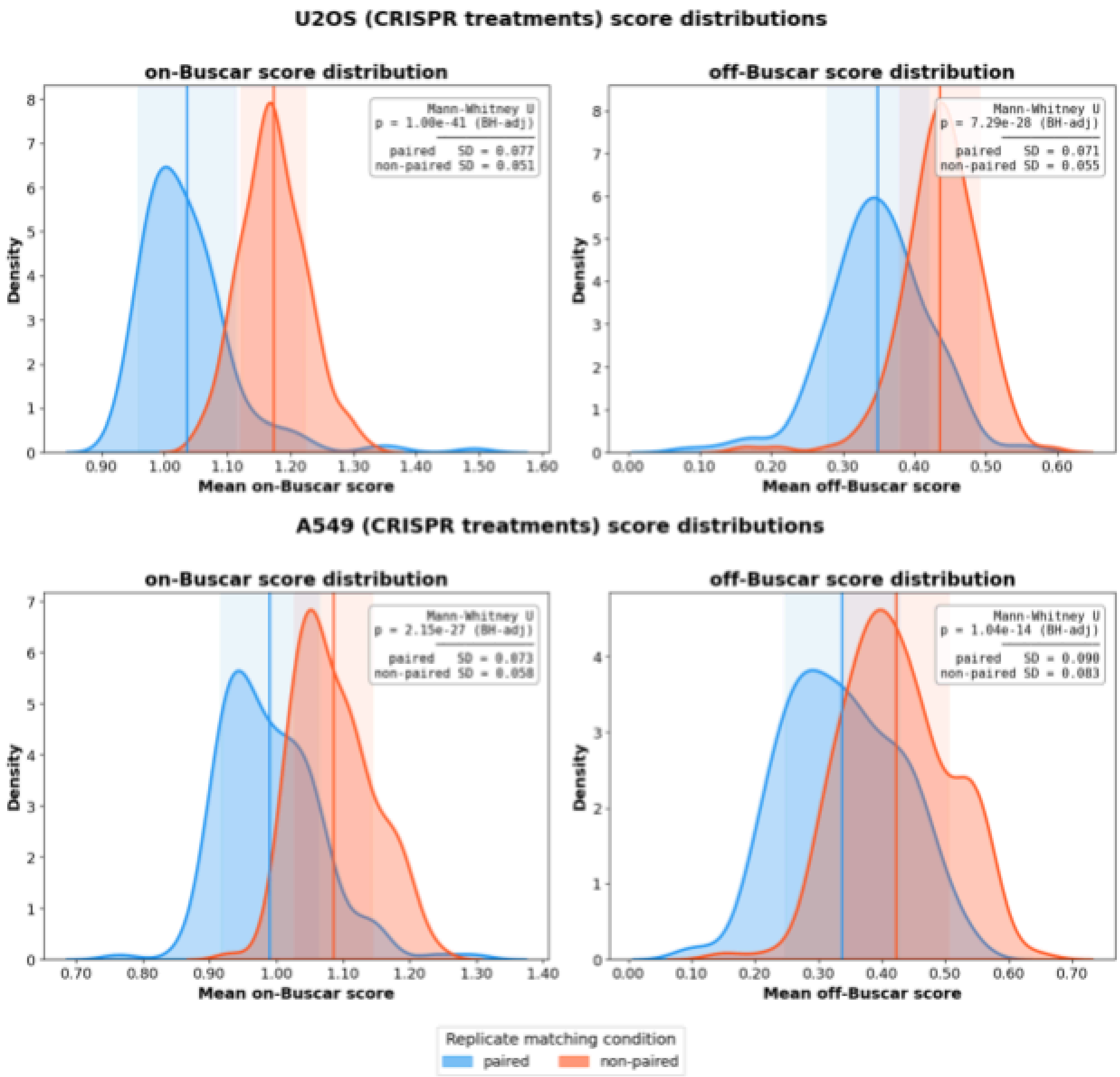

**Supplementary Figure 10. Buscar replicate analysis score distributions for CRISPR-Cas9 perturbations across U2OS and A549 cell lines.**

Each panel displays kernel density estimates of mean on-Buscar scores (left) and off-Buscar scores (right) per treatment, computed across 10 iterations in which every plate serves once as the reference. Blue distributions represent paired comparisons. Orange distributions represent non-paired comparisons. Vertical lines indicate the mean of each distribution, and shaded regions denote ±1 standard deviation. Statistical significance between paired and non-paired distributions was assessed using the Mann-Whitney U test with Benjamini-Hochberg (BH) multiple testing correction. Results for small-molecule compound treatments in U2OS (top) and A549 (bottom) cells.
